# The *Aspergillus nidulans* transcription factor SclB governs the transition from vegetative to asexual development

**DOI:** 10.1101/2025.07.23.666256

**Authors:** Emmanouil Bastakis, Rebekka Harting, Alexandra Nagel, Tanja Lienard, Gabriele Heinrich, Verena Grosse, Nicole Scheiter, Gerhard H. Braus

**Affiliations:** Department of Molecular Microbiology and Genetics, Institute for Microbiology and Genetics, University of Göttingen, Göttingen, Germany

**Author notes:** corresponding author (GHB).

**Keywords:** Transcription factors, asexual development, *Aspergillus nidulans*, *Verticillium dahliae*, *sclB*, *blrA*, *ppoC*, *veA*, *velB*

## Abstract

Asexual reproduction in filamentous fungi is a common, efficient and fast differentiation process, for producing large numbers of asexual spores (conidia), which can be distributed through the air to colonize new environments. The whole process is tightly controlled by specific regulatory proteins. Among those major regulators is the zinc-finger domain protein of SclB (Sclerotia like B), known to influence various aspects of asexual growth and secondary metabolism in *Aspergillus nidulans* as well as other filamentous fungi. Two different growth conditions of *A. nidulans* were compared to obtain a mechanistic overview for the role of SclB, mainly during the transition of the fungus from vegetative to asexual growth. Chromatin immunoprecipitation was coupled with next generation sequencing (ChIP-seq) and combined with transcriptomic analyses (RNA-seq). SclB coordinates this developmental shift mainly by controlling the expression of genes encoding for few, however, prominent regulators of conidiation. They include the transcription factors BrlA, VelB and SclB and the pheromone oxygenase PpoC. Association of SclB to promoter regions requires the newly identified SclB response element (SRE) with a nine base-pair DNA motif. Scl2 is the corresponding protein in the fungal plant pathogen *Verticillium dahliae* and partially complements the Δ*sclB A. nidulans* asexual deficiency. This supports a conserved function of this regulator among different fungal species. In summary, SclB coordinates transition from vegetative growth to asexual reproduction in *A. nidulans* through *in vivo* transcriptional control over genes coding for established players of conidia formation.

**Importance:** Fungi constantly adapt to environmental changes in their various habitats. Asexual spore formation allows to quickly leave an unfriendly habitat through dispersal into the air. The asexual developmental program of fungi ensures large number of spores, in a short period of time and in energetically efficient manner. SclB transcription factor is a key regulator of asexual growth and secondary metabolism in numerous fungal species. The mechanism through which SclB orchestrates the transition of the *Aspergillus nidulans* filamentous fungus from the vegetative to the asexual growth was revealed. This regulator directly controls *in vivo* itself as well as expression of master genes for the asexual program such as *brlA* for transcriptional control or *ppoC* for pheromone production. This study enhances the molecular understanding, how fungal asexual differentiation is initiated and coordinated, which supports the development of better strategies to control fungal pathogens, improving human health, safety and crop management.

## Introduction

Conidia are the final products of the asexual developmental program in many filamentous fungi. From the stage of mycelia (undifferentiated hyphae) till conidia production, the fungus is undertaking a variety of structural and conformational changes (1). These changes are controlled by regulatory transcription factors (TFs), which constitute the end line receivers of signals generated from external environmental inputs or internal processes (2). TFs convert those signals into specific gene expression changes leading ultimately into specific proteins necessary for the growth under specific conditions.

The asexual developmental program consists of multiple transcriptional regulatory levels and interactions. The central player for the onset of asexual sporulation is the transcriptional activator BrlA (bristle A) (3). SfgA (suppressor of *fluG*), VosA (viability of spores A) and NsdD (never in sexual development D) are TFs that occupy the *brlA* promoter and repress its expression (3–5). Under asexual conditions the FluG (fluffy G) regulator attenuates the effects of these three repressors. That leaves the promoter of *brlA* free to interact with the so-called (fluffy low brlA) FlbB, FlbC and FlbD activator proteins (6, 7). The regulators contributing to the induction of *brlA*, are called upstream developmental activators (UDAs) (6). BrlA controls genes necessary for the completion of sporulation, e.g. the formation of sporogenus phialides (*<u>abaA</u>*) (8) and the structuring and maturation of conidia (WetA) (9). AbaA (abacus A) controls the expression of *brlA*, *abaA* and *wetA* (*wet-white A*), and thereby influences the progress and completion of asexual development (8). BrlA, AbaA and WetA are key control elements of the so-called central developmental pathway (CDP) for the initiation of sporogenesis, maturation and finally the survival of conidia as asexual spores to be released into the air (10).

The velvet domain family is a fungal specific group of transcriptional regulators with central roles in development, which link differentiation to the appropriate specific secondary metabolism in *A. nidulans* as well as in other fungi (1). The velvet domain DNA binding protein associates *in vivo* to promoter regions and has the same fold as mammalian NF-Kappa (11). The ability of velvet domain proteins to form homo- or hetero-dimers provides a complex regulatory system of distinct homo- or heterodimers (1, 12). VelB is an inducer of asexual growth, since its deletion produces less conidia and shows reduced expression of the asexual regulators *brlA*, *abaA* and *vosA* (13). VeA, VelC and VosA have various prominent roles during sexual development but also operate as repressors of asexual growth (13–15).

Pheromones are essential for the reproduction of filamentous fungi. The proportion of three psi (precocious sexual inducer) factors, psiA, psiB and psiC, in *A. nidulans*, defines the shift between sexual and asexual development (1). Specifically, the dioxygenase PpoC, in charge for the synthesis of psiBβ, is an inducer of conidiation (1, 16).

Fungal zinc cluster proteins, include a great number of TFs characterized by a cysteine residue linked to two zinc atoms (17). The *A. nidulans* Sclerotia like B (SclB) zinc cluster protein has been associated with the regulation of asexual development and secondary metabolism, but also as a direct repressive target of VosA (11, 18). More recently, SclB was also shown to be involved in the coordination of secondary metabolism in fungal-fungal cocultivation (19). The implication of the protein in asexual development was originally shown in *Aspergillus niger* (20), where in a UV mutagenesis screen, two separate mutants were isolated, *scl-1* and *scl-2*. Both showed severe defects regarding asexual sporulation alongside with the formation of sclerotia-like structures during growth on plates.

Here we show that *A. nidulans* SclB is specifically important for the transition from vegetative to asexual growth. It directly affects the expression of genes coding for master regulators of asexual growth such as the transcription factors BrlA, VelB and SclB or the pheromone enzyme PpoC. *V. dahliae* Scl2 can partially rescue the strong phenotype of *A. nidulans* Δ*sclB*, which supports a conserved function of SclB among different fungal species.

## Results

### The asexual master regulator SclB has distinct *in vivo* binding and gene expression profiles across the fungal genome during vegetative compared to asexual growth

During vegetative growth SclB already controls the expression of genes coding for proteins with prominent roles during subsequent asexual development (18). Currently it is unknown whether this regulation is based on direct *in vivo* association of SclB to the promoters of these genes or whether additional intermediate regulators are involved. ChIP-seq studies were performed in combination with a transcriptomic RNA-seq gene expression analysis to examine whether this transcriptional control is based on a direct SclB interaction.

ChIP-seqs with the GFP-SclB strain were executed with mycelia grown either vegetatively for 20 h (hereafter Veg) or just after that transferred to solid medium under asexual growth conditions for another 3 h (hereafter Asex). Among the different biological replicates for each ChIP-seq, around 4000 unique gene locus IDs, were identified to be targeted by SclB in both conditions (**Fig. 1A and Fig. S1A**). In both experiments, SclB was associated *in vivo* with promoter regions spanning up to 3 kb from the transcriptional start site (TSS). Regions of the genome with a statistically significant (p<0.05) enrichment (fold enrichment ≥ 2) of mapped read sequences, as derived from GFP-SclB samples versus wildtype (hereafter Wt) (negative control/background signal), are named as peaks and indicated positions where SclB association with these regions occurred.

**FIG 1:**
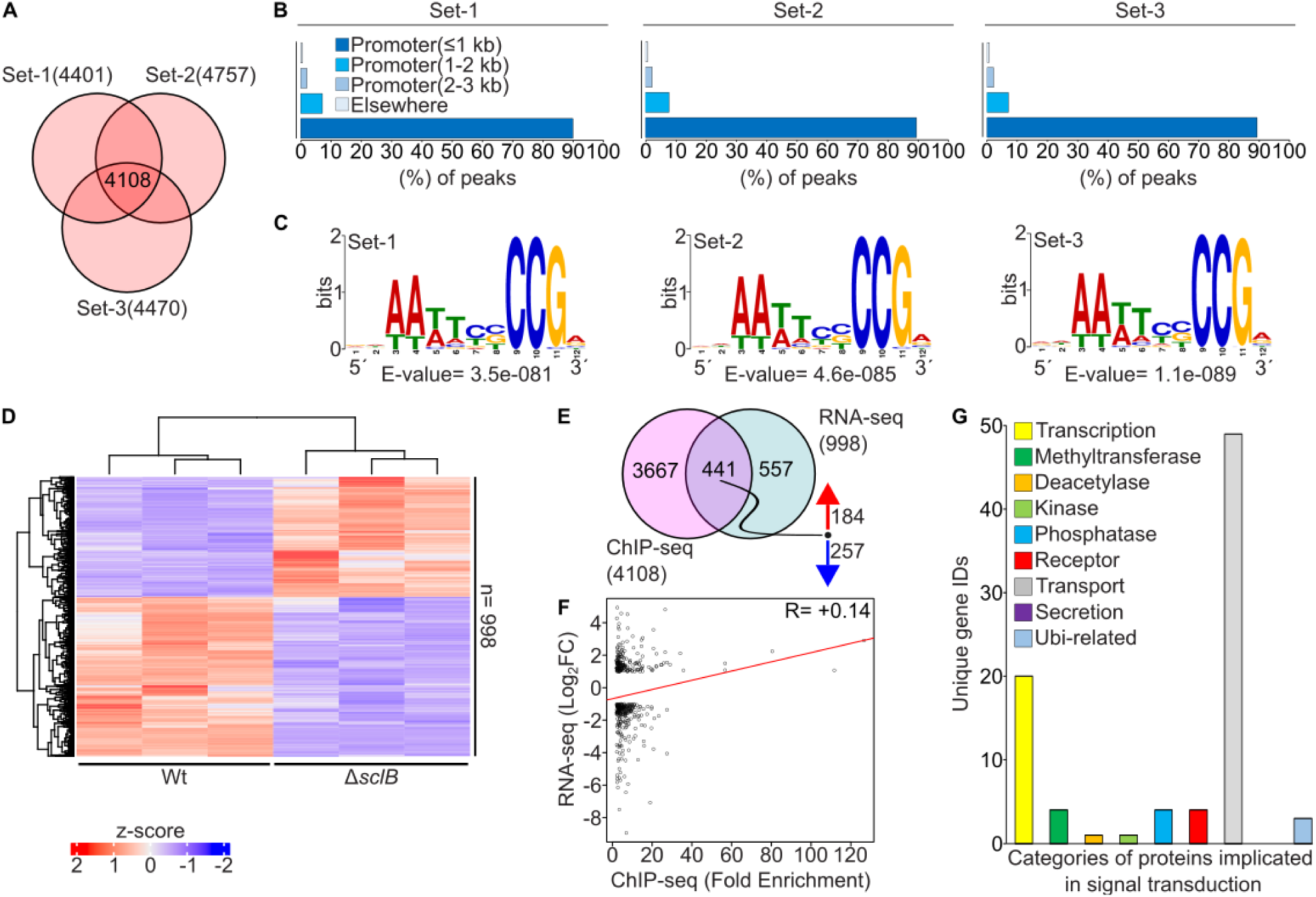
*In vivo* binding landscape of the GFP-SclB transcriptional regulator during vegetative fungal growth of *A. nidulans*. (**A**) The Venn diagram presents the overlap of three independent sets of ChIP-seq analysis for GFP-SclB versus wildtype (hereafter Wt). ChIP-seq was performed with mycelia growing for 20 h under vegetative growth conditions, (hereafter Veg). In sum 4108 unique gene locus IDs, found to be associated with peaks (cut offs: p < 0.05 and fold enrichment (F.E.) ≥ 2.0) located in up to 3 kb promoter regions, discovered simultaneously in all three independent sets of the ChIP-seq analysis. (**B**) Bar diagrams depict the distribution of the statistically significant ChIP-seq peaks over different genetic elements for each of the three independent sets of analysis. For all three sets, approximately 90 % of the identified peaks are located within promoter regions of genes up to 1 kb from the transcription start site (TSS). (**C**) Logos depict the top ranked *de novo* motifs, as discovered by using the MEME-ChIP web tool. An input set of 150 sequences consisting each of a length 100 bp was used for each independent set of analysis. All of these sequences were located below the summits of the top-ranked 150 ChIP-seq peaks for each set. The summits of all of these peaks were located up to 3 kb promoter regions. (**D)** Heatmap shows the differential expression of genes (as z-score values) from fungal mRNAs of Δ*sclB* and Wt strains, derived from mycelia growing under Veg conditions. The total number of genes found to be differentially expressed (n) under the cut offs reflect p < 0.05 and -1 ≤ log_2_FC(Fold Change) ≥ 1. The RNA-seq was performed with three independent biological replicates of Δ*sclB* and Wt strains, respectively. (**E**) Venn diagram showing the overlap among the highly reproducible targets genes from the ChIP-seq of GFP-SclB (**A**) with the DEG from the RNA-seq of Δ*sclB* (**B**). Both experiments performed with mycelia growing under Veg growth conditions. The corresponding cut offs were for the ChIP-seq: p < 0.05 and fold enrichment (F.E.) ≥ 2.0 and for the RNA-seq: p < 0.05 and -1 ≤ log_2_FC(Fold Change) ≥ 1. The numbers beside the red and blue arrows indicate upregulated or down-regulated genes from the total genes of the overlap. This set of 441 locus IDs reflects the direct *in vivo* targets of GFP-SclB for mycelia growing under Veg conditions. (**F**) Scatter plot showing the Pearson correlation coefficient between the peak’s fold enrichment from the ChIP-seq in (**A**) with the gene expression as of Log_2_FC from the RNA-seq in (**B**), under the same Veg growth. For this plot only the 441 direct target genes found in the overlap of ChIP-seq with the RNA-seq in (**E**) were used. The R= +0.14 inside the plot indicates the positive correlation between the specific NGS data sets tested. (**G**) Bar chart depicting the number of genes from the overlap (441 gene locus IDs) between the ChIP-seq and RNA-seq (as shown in **E**) belonging to categories of genes encoding proteins, known to be implicated in signal transduction processes. The number for each category was defined after performing a manual search of the set of 441 gene locus.

Subsequently, it was examined how the unique peaks are distributed over different genetic elements along the fungal genome. It was found that close to 90 % of the identified peaks, for vegetative or asexual ChIP-seqs, where located in promoter regions of genes that are extended up to 1 kb from the TSS (**FIG. 1B and Fig. S1B**). The DNA binding motif, through which SclB association occurs was elusive. Sequences of 100 bp located directly below the summit of the 150 top scored peaks (based on the fold enrichment) were used to discover this motif for each independent set of the ChIP-seq analysis. A nine base pairs long DNA motif (consensus: 5’-AATTCCCCG-3’) with consistent composition among the different sets was discovered (**Fig. 1C and Fig. S1C**). This motif is recognized by SclB and was accordingly named SclB response element (SRE).

The *in vivo* association of a TF does not always mean immediate transcriptional regulation of the nearby genes, because coregulation with other regulators is required or the change in the expression of gene(s) takes place at a different developmental point or condition (21). Hence, the binding events of SclB were identified, which are truly leading to direct transcriptional regulation of genes located in close proximity to the corresponding peaks. Therefore, genome-wide gene expression analyses by RNA-seq assays were performed under the same vegetative and asexual inducing conditions (Veg/Asex) with the ChIP-seqs, by comparing Δ*sclB* and Wt strain cultures. The principal component analysis among the different replicates of each of the sample groups, showed a clear separation and clustering between both RNA-seqs (**FIG S2**). A total number of 998 genes for the Veg and 609 for the Asex were found to be differentially expressed (cut offs: p < 0.05 and -1≤log_2_FC≥1) when comparing Δ*sclB* and Wt samples (**Fig. 1D and Fig. S1D**). Moreover, the visual representation of these differentially expressed genes (here after DEGs), as indicated in the corresponding heatmap, illustrated a distinct expression profile (presented as z-score values) of those genes among the tested growth conditions.

We then set to define, by the overlapping of these two data sets, the so-called direct *in vivo* target genes of SclB. A set of in total 441 unique gene locus IDs was found to be simultaneously differentially expressed and targeted by GFP-SclB to their promoter regions of up to 3 kb from the TSS *in vivo* (**Fig. 1E**) for vegetatively growing mycelia. However, the number of the direct SclB target genes was reduced almost in half (241) for mycelia induced for asexual development (**Fig. S1E**). Comparisons between SclB up- and downregulated target genes revealed that SclB mostly operates mostly as an inducer during Veg and as a repressor during Asex growth conditions, respectively. A calculated Pearson correlation coefficient among the corresponding ChIP-seqs and RNA-seqs revealed a clear linear positive correlation (+0.14) for vegetative (**Fig. 1F**), in contrary to a slightly negative correlation (−0.06) during asexual growth conditions, respectively (**Fig. S1F**).

SclB controls various gene regulatory networks (18). The sets of 441 (Veg) and 241 (Asex) direct target genes of SclB were manually searched for candidates encoding proteins with functions in signal transduction. Under both conditions the TF affected two overrepresented categories, which correspond to genes encoding proteins associated either to cellular transport or are related to transcription (**Fig 1G and Fig S1G**). Further analyses with a focus on biological processes (BP) of the same sets for statistically significant enrichment of GO (Gene Ontology) revealed that most other overrepresented categories were associated with processes related to fungal primary or secondary metabolism (**Fig. S3A and S3B**).

The *in vivo* binding profiles and the expression data from vegetative and asexual growth were further compared for overlaps and differences The high positive Pearson correlation coefficient (+0.94) supports that the *in vivo* targets of SclB are largely maintained between vegetative and sexual growth (**Fig. 2A and 2B**). The two transcriptomic data sets further corroborate this finding with a positive (+0.5) correlation (**Fig. 2C**).

**FIG 2:**
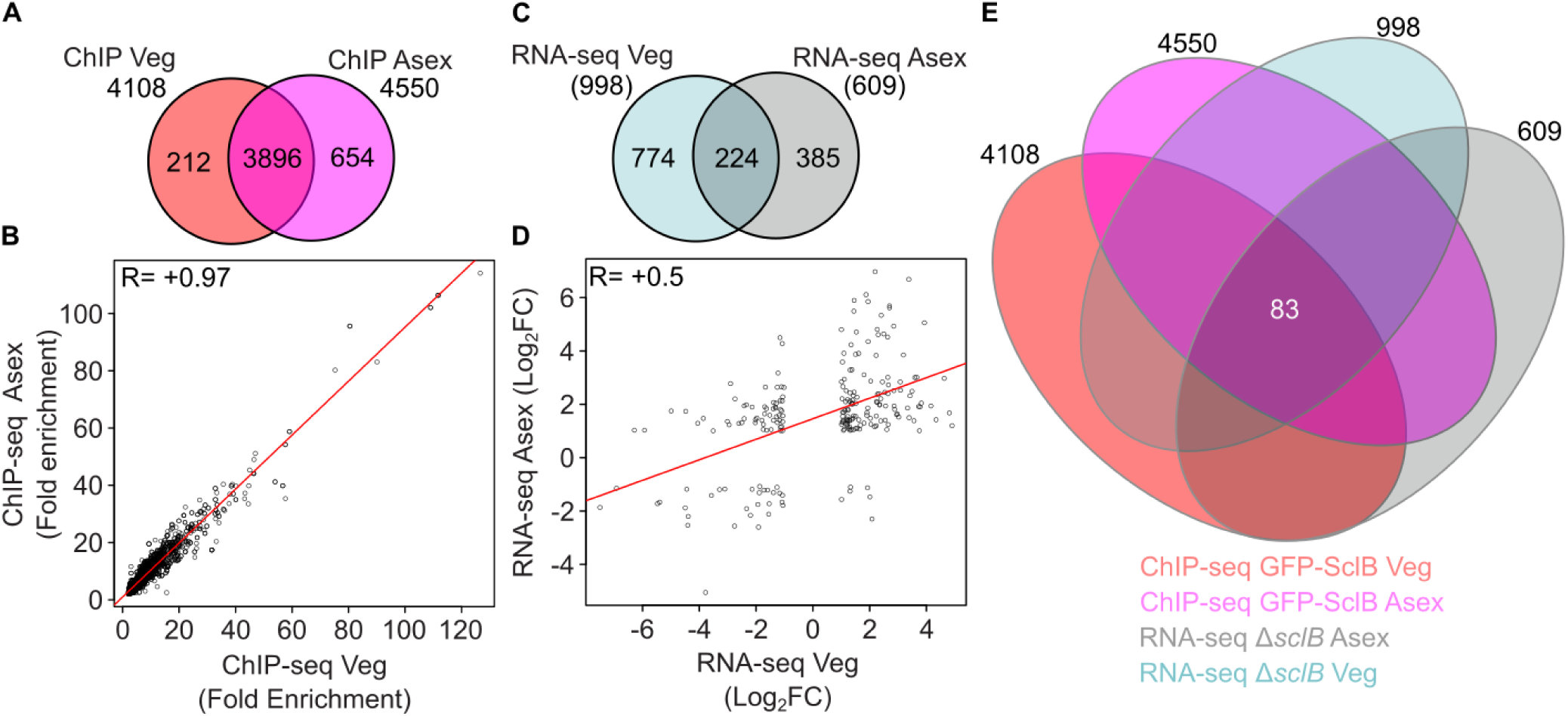
SclB directly targets 83 *A. nidulans* genes during different parts of the life cycle. (**A**) Venn diagram is illustrating the overlap of 3896 gene locus IDs between the ChIP-seq experiments conducted under the two different growth conditions of Veg (as presented in FIG 1A) and Asex (as presented in FIG 2A) growth as well. For both of the ChIP-seqs a prior filtration was applied for the identified peaks under the cut offs: p < 0.05 and fold enrichment (F.E.) ≥ 2.0, the peak must be located up to 3 kb upstream from the TSS of every gene locus. (**B**) Scatter plot illustrating a Pearson correlation coefficient, between the fold enrichment of peaks derived from the ChIP-seq of the Veg growing mycelia and the ChIP-seq with Asex growing mycelia. For both of the chip-seq experiments they were applied the filtering under the conditions: p < 0.05 and fold enrichment (F.E.) ≥ 2.0. For both of the ChIP-seq experiments, it was used the set of locus IDs of genes that showing reproducibly (in all three-independent sets of each analysis) peaks to promoter regions up to 3 kb to their promoters. The R= +0.97 at the upper left corner of the plot shows the highly positive correlation among the compared NGS data sets that were examined. (**C**) Venn diagram depicting the overlap of gene locus IDs found to be differentially expressed under both conditions examined by RNA-seq (as presented in Fig 1D and Fig 2D). Mycelia derived from Δ*sclB* and Wt strains, under Veg and Asex growth conditions correspondingly. Each RNA-seq is consisting of three biological replicates of the Δ*sclB* and another three biological replicates for the Wt strain. Both of the RNA-seq experiments were subjected to the cut off: p < 0.05 and -1 ≤ log_2_FC(Fold Change) ≥ 1. (**D**) Scatter plot presenting a Pearson correlation coefficient, among the expressions (log_2_FC) of gene found to be differentially expressed in the RNA-seq with the Veg grown mycelia and with the Asex grown mycelia. Both RNA-seq experiments were subjected to cut offs: p < 0.05 and -1 ≤ log_2_FC(Fold Change) ≥ 1. The R= +0.5 at the upper left corner of the scatter plot indicates a strong positive correlation among the DEGs among the different RNA-seq data sets. (**E**) Multi-Venn diagram highlighting a total number of 83 gene locus IDs found to be identified in both of the ChIP-seqs and both of the RNA-seqs performed under Veg and Asex growth conditions.

In contrast, the overlap of the differentially expressed genes (224 DEGs) between the two RNA-seqs was found to be severely reduced at roughly one fifth from the Veg and one third for the Asex data set, respectively (**Fig. 2D**). Genes, which are conserved direct targets of SclB under vegetative as well as asexual growth conditions were identified by a Venn diagram between all four NGS datasets. A set of 83 genes could be identified as SclB target genes that remained associated with the transcription factor under all tested conditions (**Fig. 2E**). The comparison of the expression patterns of these 83 genes revealed that 57 candidates maintained the same expression pattern during vegetative or asexual growth (**Fig. S4**). In contrast, 26 genes had an opposite expression profile after the transition to the asexual phase (**Fig. S4**). In sum, these results suggest that the *in vivo* affinity of SclB to promoters of a particular set of 83 genes (mostly related with metabolism, cellular transport and transcription), is largely conserved during the shift from vegetative to asexual growth.

### SclB mediated transcriptional control of *brlA*, *ppoC* and *sclB* orchestrates the transition from fungal vegetative to asexual growth *in vivo*

SclB influences the expression of genes encoding major key regulators of asexual development (18) in mycelia grown either 24 h vegetatively or during final stages of asexual development (24 h post induction). The direct influence of SclB on the expression of core asexual regulators was examined, specifically during the transition from vegetative to asexual growth, by comparisons with a list of established asexual regulators published by Krijgsheld et al., (22). This list consists of 35 genes, to which we added one more, *sclB* (AN0585) itself, published and characterized at a later time by Thieme et al., 2018. The set of 36 genes in total of established asexual regulators (hereafter asexual MRs), was used for the comparisons.

Different overlaps with the list of the asexual MRs and the Veg and Asex NGS data sets of SclB were performed. Independently, from the growth condition, there was always a very small set of genes, *sclB*, *ppoC* and *brlA,* from known asexual MRs, that was found to be directly regulated by SclB (**Fig. 3A and 3D**). However, the number of genes encoding asexual MRs was larger when the overlaps performed with genes only appeared in the ChIP-seq lists (**Fig. 3B and 3E**). When overlaps were performed with the lists of DEGs from the RNA-seqs, it was again a small set of genes emerged as common loci IDs (*brlA*, *sclB, ppoC*, *sclB*, *flbD*, *flbC*, *ppoB* and *rodA*). In summary, SclB shows a rather strict selective behavior during the transition of mycelia from vegetative to asexual growth, concerning the transcriptional control of genes encoding for key regulators of asexual development. Genes coding for the prominent master regulators, *brlA*, *ppoC* and *sclB* itself, are not only targeted by SclB *in vivo* but also become differentially expressed by it.

**FIG 3:**
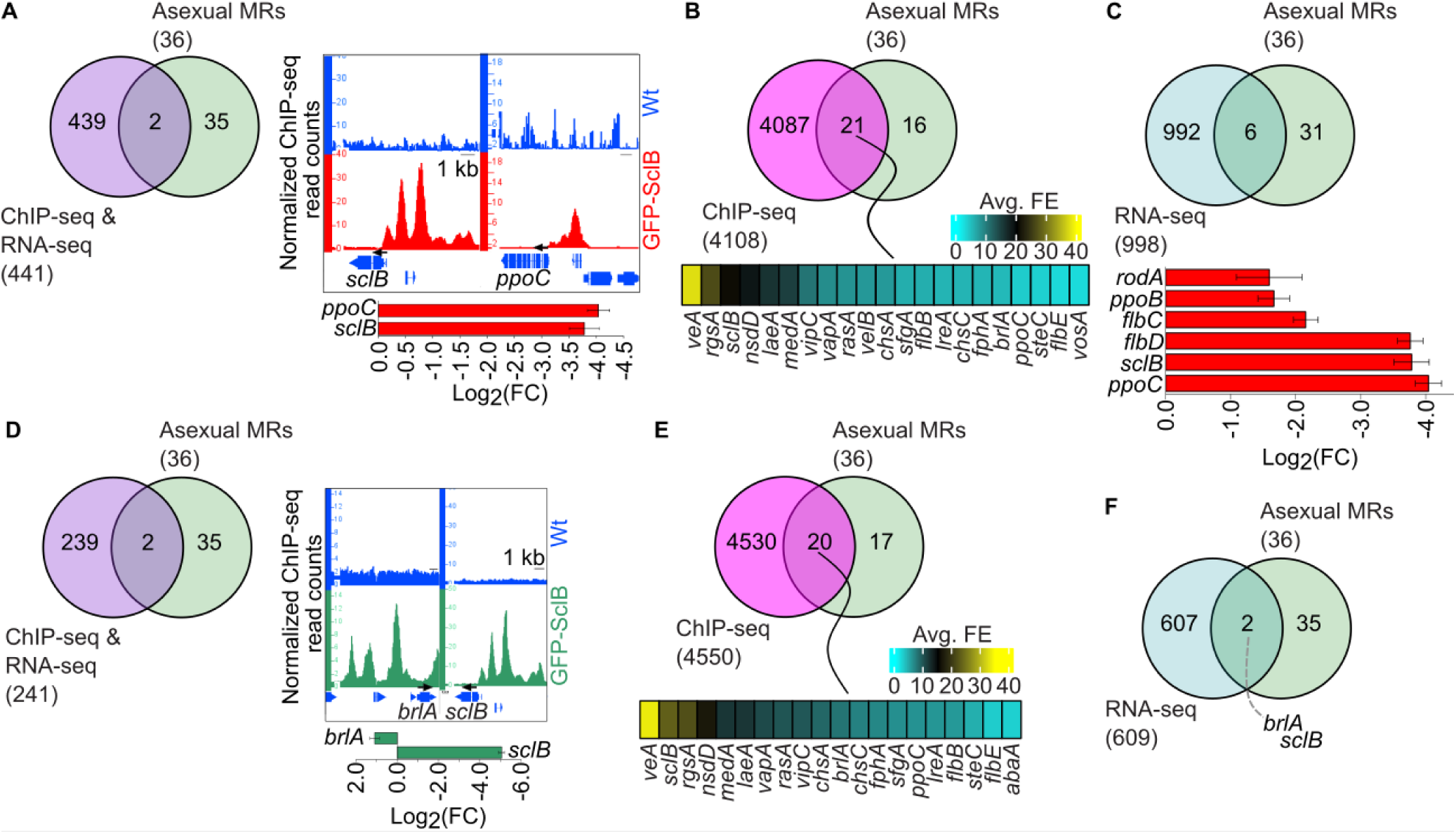
*A. nidulans* SclB exerts direct transcriptional control on its own expression together with expression of the regulatory gene *brlA* and the *ppoC* gene encoding an enzyme for oxylipin formation. Venn diagrams are illustrating the overlap of the direct target genes, as found from the ChIP-seq (cut offs: p < 0.05 and fold enrichment (F.E.) ≥ 2.0) and the RNA-seq (cut offs: p < 0.05 and -1 ≤ log_2_FC(Fold Change) ≥ 1 experiments, from mycelia growing under (**A**) Veg or (**D**) Asex conditions, with the list of 36 asexual MRs genes. The right parts in panels (**A**) and (**D**) show screen shots from the Integrative Genome Browser (hereafter IGB) where peaks (*in vivo* binding positions) of the GFP-SclB to the promoters of genes from the overlaps are shown. The bar plots, below the IGB screen shots depict the expression of genes as found in the RNA-seq, performed with RNAs derived from mycelia of Wt and the Δ*sclB* strains growing for either under Veg or Asex conditions. Venn diagrams presenting the overlap between the ChIP-seq strong targets, performed with mycelia derived from either (**B**) Veg or (**E**) Asex growth with the list of asexual MRs genes. The heat maps, underneath the Venn diagrams, depict the strength (as fold enrichment color key) for the *in vivo* binding of GFP-SclB to the promoters of the 21 genes (in **B**) and the 20 (in **E**) genes found in the corresponding overlaps. Venn diagrams showing overlaps between the strongly differentially expressed genes, as derived from the RNA-seq, with RNAs from mycelia from Wt and Δ*sclB* strains growing under (**C**) Veg or (**F**) Asex conditions, with the list of the asexual MRs genes. The bar plot below of the Venn in (**C**) illustrates the expression (as of log_2_FC) of the 6 genes found in the corresponding overlap.

### Members of the emericellamides gene cluster are direct targets of SclB during vegetative and asexual growth

The zinc-finger transcriptional regulator SclB has been strongly associated with the regulation of secondary metabolism in several Aspergilli. *A. nidulans* SclB is inducing the biosynthesis of emericellamides (23) during vegetative growth and austinol and dehydroaustinol under asexual growth conditions (18), the corresponding *scl2* orthologue of *A. niger* is controlling the biosynthesis of indoloterpenes/ aurasperones (20). It was further investigated, whether there is a direct transcriptional SclB control in *A. nidulans* towards members of the *eas* and *aus* biosynthetic gene clusters (BGCs) during the transition from vegetative growth to asexual development. These BGCs are responsible for the biosynthesis of emericellamides, austinol and dehydroaustinol correspondingly. The ChIP-seq data from both experimental conditions were searched for the presence of peaks on promoter regions of genes of the *eas* and *aus* cluster. In the case of the *aus* BGC, one clear peak was discovered under both experimental conditions (**Fig. 4A**). However, examination of the corresponding transcriptomic data showed that the *aus* genes, which were differentially expressed were not in close proximity to the SclB location *in vivo* (**Fig. 4C and 4A**). However, strong ChIP-seq peaks were identified under both conditions at the promoters of all four genes (*easA-D*) coding for members of the *eas* BGC for emericellamides production (**Fig. 4B**). The expression of the *eas* genes during Asex growth was found to be repressed by SclB (**Fig. 4D**). In contrast, the expression of the same genes was found to be induced by SclB during vegetative growth (**Fig. 4F**). This corroborates that SclB is directly associated with promoters of genes coding for BGCs *in vivo*. Furthermore the regulatory role of SclB changes during the transition from vegetative growth to asexual development, from being inductive to becoming repressive concerning the *eas* genes in particular.

**FIG 4:**
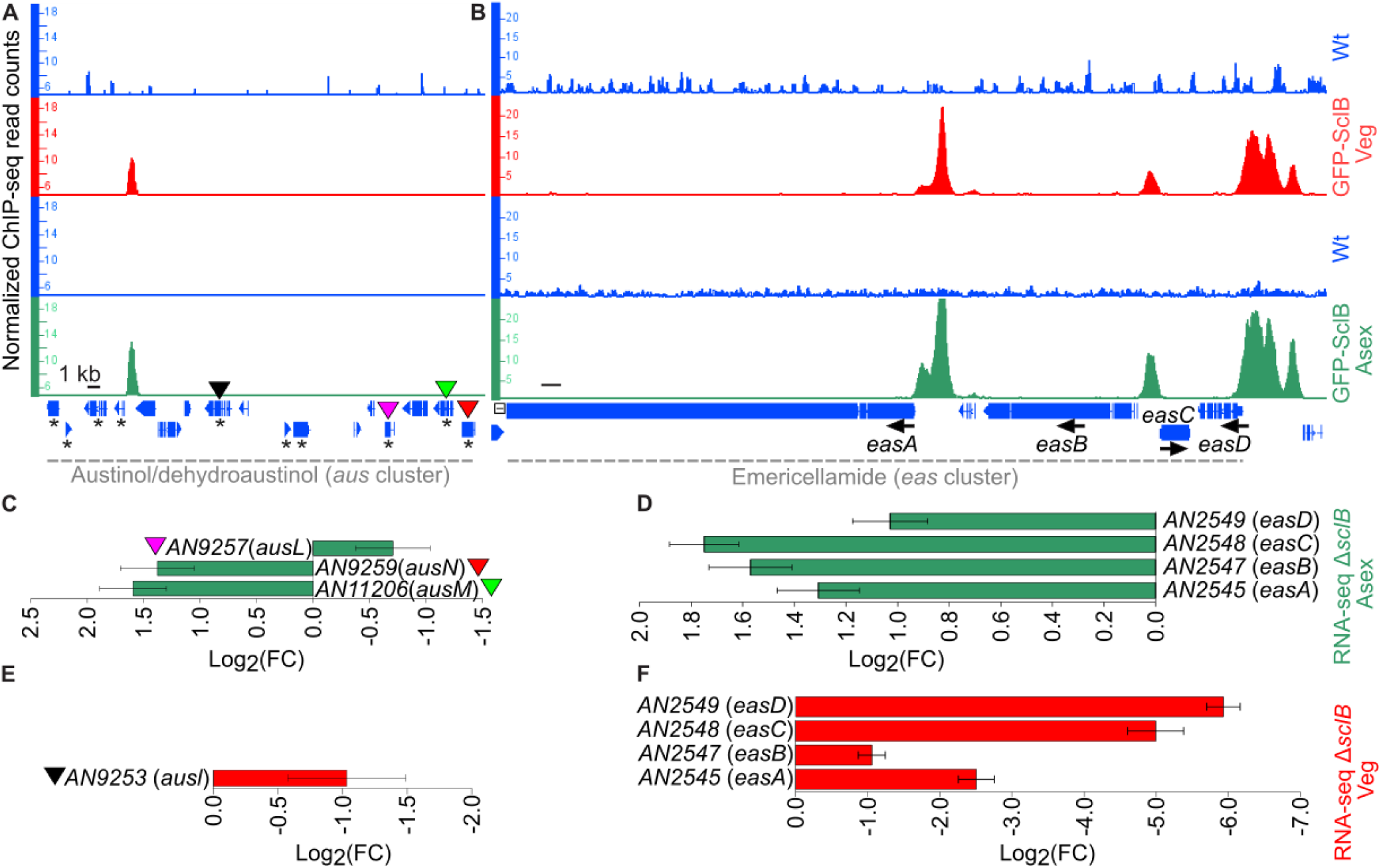
SclB regulates the expression of *A. nidulans* genes of the *aus* and *eas* secondary metabolite gene clusters. Snapshots from the IGB depicting the *in vivo* binding of SclB (ChIP-seq peaks) on promoters of genes of the *aus* (**A**) and *eas* (**B**) gene clusters. Peaks appeared in the red tracks illustrating the *in vivo* binding of SclB from mycelia growing under Veg. Peaks presented in green tracks derived from mycelia growing on Asex conditions. Blue tracks show in each case the corresponding negative control (background signal) of the ChIP-seq along the corresponding gene clusters. Green bar charts, are illustrating the expression (as log_2_FC) of genes of the *aus* (**C**) and the *eas* (**D**) clusters, found to be differentially expressed (p < 0.05) in the RNA-seq data generated by mycelia of the Δ*sclB* and Wt strain, growing on under Asex growth conditions. In panel (**C**) the colored triangles show the correspondence of discovered DEGs in the RNA-seq with their specific locus in the ChIP-seq data of panel (**A**). The red bar charts showing the expression (as log_2_FC) of genes of the *aus* (**E**) and the *eas* (**F**) clusters, found to be differentially expressed (p < 0.05) in the RNA-seq data generated by mycelia of the Δ*sclB* and Wt strain, growing under Veg conditions.

### *veA* and *velB* are direct target genes of SclB

The *A. nidulans* velvet-domain proteins and their orthologs in other species, have been associated with a great variety of aspects of fungal development (1). Each of the four velvet family members (VeA, VelB, VelC and VosA) has been linked to distinct processes through the life cycle of the fungus. VosA was found to directly associate to *sclB in vivo* (11). A later study linked the VosA binding to the transcriptional regulation of the *sclB* gene (18). It appeared that VosA is a repressor of *sclB*, at least in asexually grown fungal colonies. We hypothesized that the regulation among VosA and SclB can be i) mutual and ii) possibly SclB can transcriptionally control other velvet genes besides *vosA*. To test this, SclB ChIP-seq data were searched for peaks nearby the promoters of the velvet genes. The promoter regions of *veA*, *velB* and *velC*, were found to be occupied by SclB at multiple positions (**Fig. 5A**). Binding signals (peaks), indicating an *in vivo* association of SclB, were found at several positions in the *vosA* promoter. However, these binding events would rather be characterized as weak and transient, especially compared to the ones found in the promoters of the other velvet genes. Notably, the strong *in vivo* association of SclB to the promoters of *veA*, *velB* and *velC* was taking place independently from the growth conditions (**Fig 5A**). Subsequently, it was examined whether these binding events could provoke changes of velvet gene expression. Investigation of the corresponding RNA-seq transcriptomic data revealed that only two out of four velvet genes, *veA* and *velB*, were repressed in Δ*sclB* compared to the Wt, when the mycelia of the fungus were grown under vegetative conditions (**Fig. 5B**). This highlights the inductive role of SclB when associated with the promoter regions of the corresponding velvet genes (**Fig. 5A**). During early asexual growth only *velB* was repressed in Δ*sclB* compared to Wt (**Fig. 5B**). These data suggest that SclB binds to the promoters of the *veA* and *velB* genes and presumably support their activation, whereas the function of the interaction of SclB to the other promoters of velvet genes remains yet elusive.

**FIG 5:**
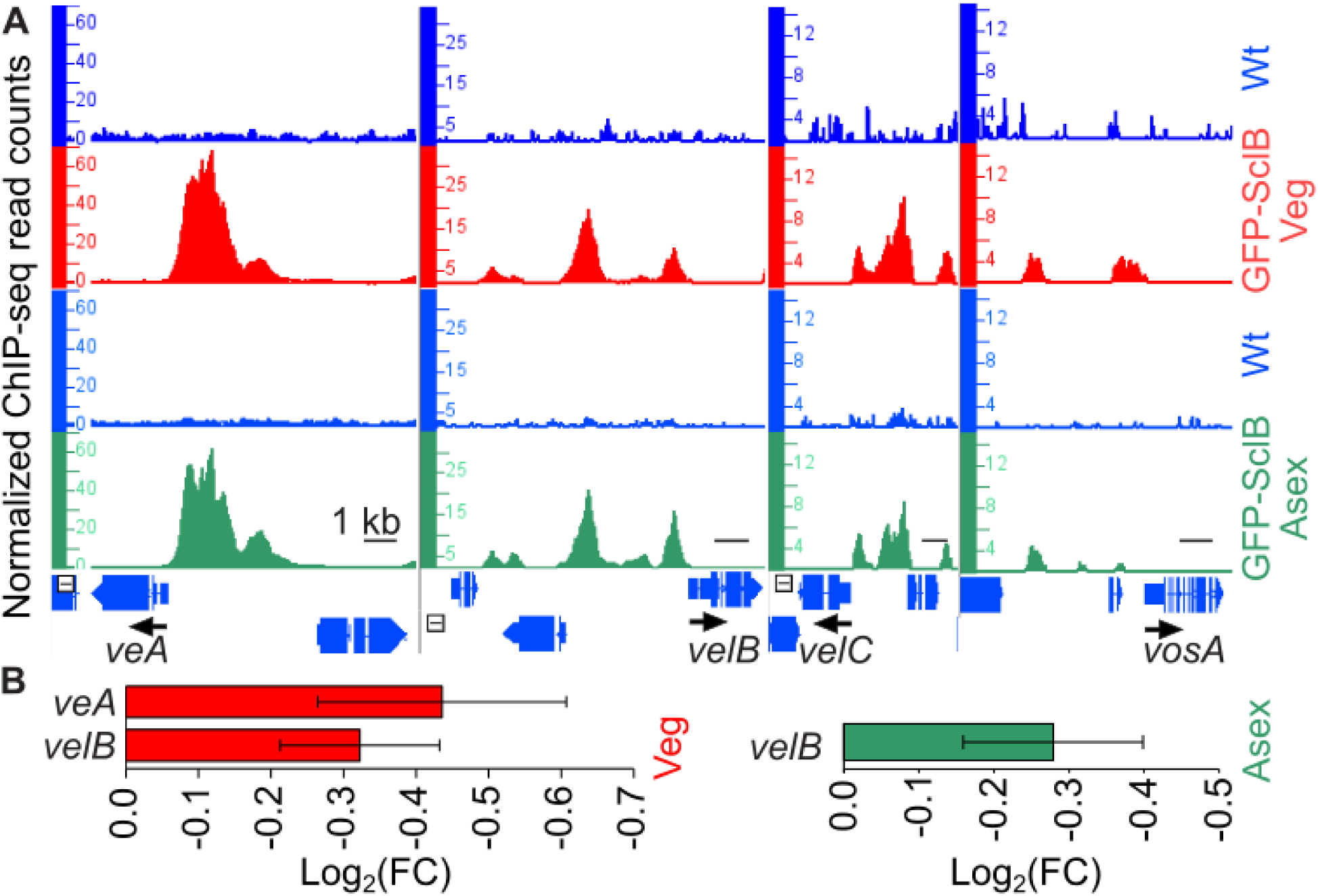
Expression of fungal *veA* and *velB* regulatory genes is induced by the *in vivo* association of SclB to their promoters. (**A**) Screen shots from the IGB depicting the *in vivo* binding of SclB (ChIP-seq peaks) to the promoters of the velvet genes (*veA*, *velB*, *velC* and *vosA*). Peaks appeared in the red tracks illustrate the *in vivo* binding of SclB in mycelia growing under Veg conditions. Peaks showed in green tracks illustrate *in vivo* binding of SclB for mycelia under Asex growth. Blue tracks show in each case the corresponding negative control (background signal) of the ChIP-seq. (**B**) Bar charts, just below the ChIP-seq snapshots are illustrating the expression of *veA* and *velB*, as found to be differentially expressed in the RNA-seq data (p < 0.05) generated by mycelia of the Δ*sclB* and Wt strains, growing either under Veg (red bars) or under Asex growth conditions (green bar).

### The *scl2* ortholog of *sclB* from the plant pathogen *Verticillium dahliae* encodes a protein with partial conserved function with SclB from *A. nidulans*

*A. nidulans* uses asexual spores to disseminate and colonize new habitats. In contrast, the plant pathogenic fungus *V*. *dahliae* produces spores when it reaches the host xylem sap to colonize the upper parts of the plant (24, 25). We wanted to investigate if SclB is conserved in another filamentous fungus that uses the production of spores in a very distinct environment only. BlastP (26) was employed using the amino acid sequence from *A. nidulans* SclB as a reference. We discovered two neighboring genes, VDAG_JR2_Chr8_g03870a and g03890a. None of the deduced amino acid sequences had any predicted domains. It was then assumed that this might be a misannotation, hence a proposed cDNA transcript of one gene was amplified and sequenced (**Fig. S7**). The deduced amino acid sequence of *Vd* Scl2 shares approximately 48% identity with *An* SclB (using MUSCLE alignment) and a Zn(2)-C6 fungal-type DNA-binding domain was predicted (IPR001138). The open reading frame of *Vd scl2* was deleted and the phenotype of the resulting strain was investigated. There were no significant differences in colony morphology or production of conidiospores compared to Wt (**Fig. S5**).

It was examined whether the putative Vd Scl2 protein might have retained some functions, thus maybe is able to complement the phenotype of the *An* Δ*sclB* strain. Therefore, *A. nidulans* strains were generated in locus expressing *Vd scl2* with or without a C-terminal GFP (green fluorescent protein), in Wt and Δ*sclB* background strains. The nuclear localization of the Vd Scl2 was subsequently tested by employing confocal live imagining microscopy in the Wt (as a negative control/background signal) and all the complementation strains where *Vd* Scl2 was tagged at the C’-terminus with GFP. All strains were transformed with the universally used RFP-H2A red nuclear fluorescent marker as a reference for nuclear signal. *Vd* Scl2-GFP either expressed in the Wt or in the Δ*sclB* strain, was always observed nuclear (**Fig. 6A**).

**FIG 6:**
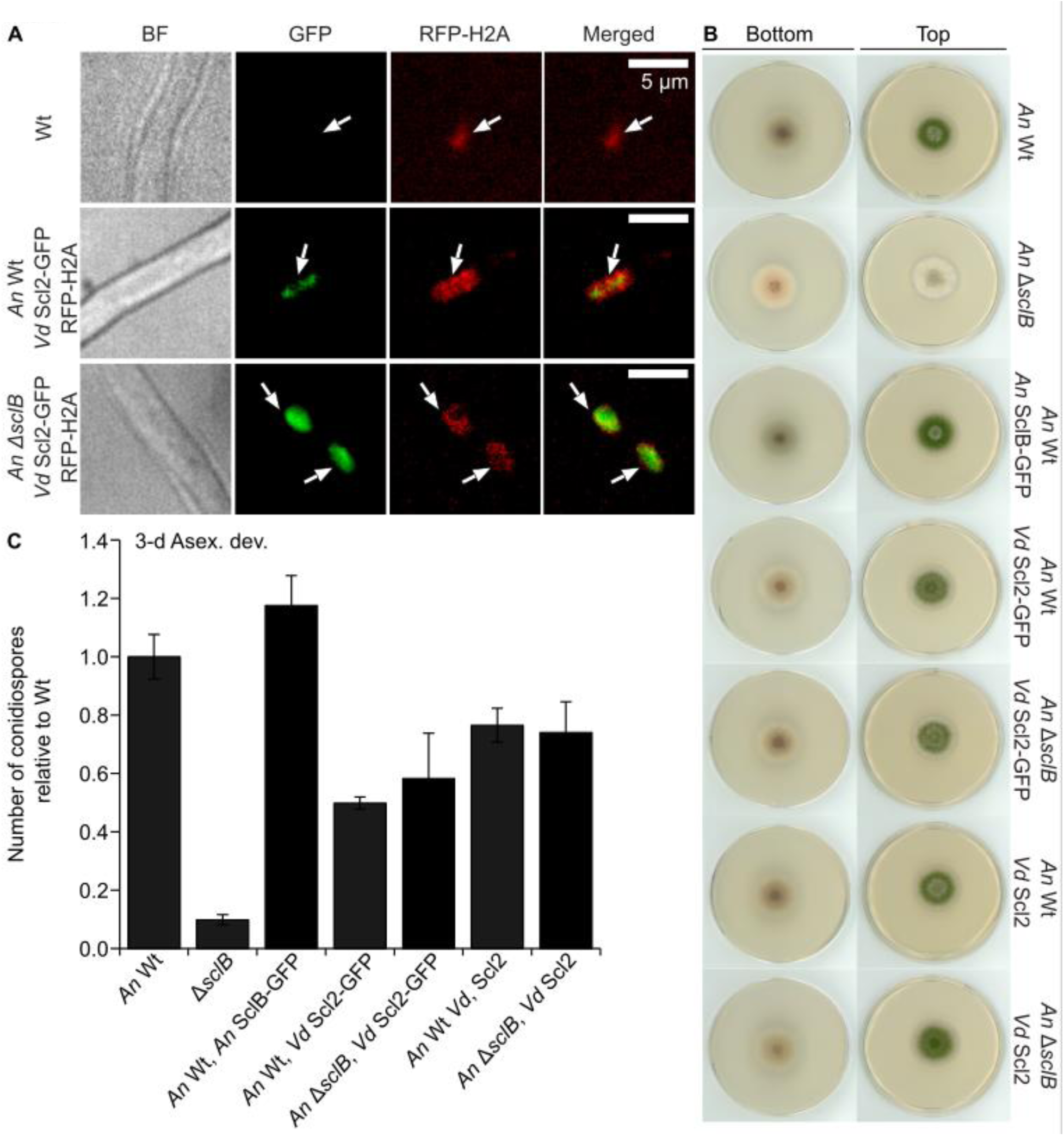
The nuclear Scl2 protein from plant pathogenic *V. dahliae* can overtake SclB regulatory functions and partially rescues the defective phenotype of the *A. nidulans* Δ*sclB* mutant strain. (**A**) Cellular localization of the *Vd* Scl2 tagged by a green fluorescence protein (GFP) from *V. dahliae* (*Vd* Scl2-GFP), while was expressing in the Wt and in the Δ*sclB* strains of *A. nidulans*. In both cases the Scl2-GFP is expressed under the native promoter of *An sclB*. The *An* Wt, the *An* Wt *An* SclB-GFP and the *An* Δ*sclB An* SclB-GFP strains, carried also a transgene expressing the nuclear marker of H2A tagged with the RFP (red fluorescent protein). For the confocal images, hyphae were growing under Veg growth conditions. (**B**) Phenotypical characterization of *A. nidulans* strains, growing under asexual growth conditions for 3 days. Each plate was initially inoculated in the middle with a total of 2000 spores. After a period of 3 days, scans were made from the upper and bottom part of the plates. (**C**) Quantification of the number of asexual spores (conidia), for all the strains presented in (**B**), after 3 days growing under asexual growth conditions (constant light and 37 °C). The final quantification results for each strain derived from a minimum of at least three replicates (plates). Every plate was initially inoculated by a total of 30.000 spores, which were then spread all over the plate by using glass beads. Results for the quantification are normalized relative to the Wt strain and are representing averages and standard deviations of at least three biological replicates.

Next, it was tested if the nuclear localized *Vd* Scl2 can rescue the phenotype of *An* Δ*sclB*. After three days of asexual growth all the strains carrying *Vd scl2* formed colonies very similar to Wt (**Fig. 6B**). Quantification of asexual spores revealed that all versions were functional. The amount of the produced conidiospores did not reach Wt levels, however, more spores were produced than in the Δ*sclB* strain (**Fig 6C**). In conclusion these data strongly support that Scl2 from *V. dahliae* as well as SclB from *A. nidulans* are both nuclear localized proteins, which are conserved to a large degree in their functions between both fungi.

## Discussion

Airborne spores produced from fungi are necessary for their propagation and survival in a variety of environments. In filamentous fungi, such as *A. nidulans*, conidiospores are the final products of the asexual developmental program. During that process several genes, coding for essential proteins, need to be regulated and coordinated in temporal and spatial manner. The SclB TF is playing a pivotal role in the regulation of the expression of genes, product of which, influence many aspects of the asexual development and secondary metabolism (18), not only in *A. nidulans* but also in other filamentous fungi as well. However, it is not clear yet how SclB applies its regulatory roles particularly during the outset of the asexual development. Here, we studied the influence of SclB at the transition of the fungus from the undifferentiated vegetative growth into the early asexual development on a genome-wide scale.

The high-throughput genome-binding experiments showed for the first time that SclB can be associated *in vivo*, via a novel nine-base-pairs DNA motif (SRE), to around ca. 4000-4550 genes, which are approximately half of the total genes in the *A. nidulans* genome, regardless the growth conditions (**Fig. 1, S1 and S3 Table**). Moreover, there is a very high correlation among these two data sets, indicating a conserved preference that SclB shows for promoters of specific genes (**Fig. 2A**). However, these binding events could lead to actual changes in the expression of only a smaller number of genes, 241 for Veg and 441 for Asex growth correspondingly (**Fig. 1, S1, and S3 Table**). This could imply further roles that SclB might have during the particular shifting, related to responses to other external or internal inputs that might accompany this transition. Future studies are necessary to further elucidate yet uncovered developmental roles of SclB in fungal life.

A major function of SclB in the developmental transition is the control of the key regulatory genes *ppoC*, *brlA*, *veA* and *velB* combined with an auto control of *sclB* itself (**Fig. 3 and 5**). The direct transcriptional autoregulation of SclB to itself, alongside with the control of the gene for the dioxygenase PpoC, are two novel discoveries of the complexity of the SclB regulatory network, that were hidden so far. Specifically, the implication of SclB in the synthesis of the oleic acid-derived psi factor psiBβ, a known inducer of asexual growth (1, 16), highlights the variety of control levels via which SclB can influence the initiation of the asexual growth. It has been shown that *brlA* expression is induced if SclB is overexpressed in mycelia growing for 24 h vegetatively (18). Moreover, it is also known that *brlA* induction starts at least 5 h post induction of asexual development (27). SclB keeps the expression of *brlA* in a low level at least up to the point of 3 h post induction. Our study supports that SclB maintains first a low *brlA* expression until the first three hours of the induction of asexual growth (**Fig. 3**). This illustrates the finding of a subtle fine tuning that SclB asserts towards BrlA as part of the developmental transition. In the future, it will be informative to examine what is the actual effect of SclB on the expression of *brlA* in later time points such 5 h and more post induction. This study has established a novel transcriptional relationship between SclB and the velvet domain regulatory genes *veA* and *velB*, coding for regulators with opposite roles in asexual growth (13–15). In fact, SclB is able to induce the expression of both velvets under vegetative conditions, but after the shift to asexual growth, its inductive action was delivered only to *velB* (**Fig. 5**). This is in line with the promoting role of VelB during the asexual growth of the fungus. The MsnA regulator of development, binds *in vivo* to the *sclB* promoter and induces gene expression for vegetative and asexual growth Bastakis et al., (28). SclB is strongly associated to multiple *msnA* promoter sites during Veg or Asex growth conditions (**Fig. S6**). However, none of these binding events led to differential expression of *msnA* (**S3 Table**). It is yet elusive, whether these direct promoter interactions might result in transcriptional control of the *msnA* promoter at different time points than the one tested by our RNA-seqs.

Chemical compounds derived by the secondary metabolism of fungi, can trigger developmental processes but also shape the interactions of the fungus with other living organisms nearby (29). SclB is affecting *in vivo* the expression of genes belonging to *eas* and *aus* gene clusters, responsible for the synthesis of emericellamides (compounds with antibiotic action) (23) and austinol/dehydroaustinol (important in sporulation) (30, 31) correspondingly. This type of regulation shows that changes in secondary metabolites, triggered by SclB, are necessary during the passage of the fungus from vegetative to the asexual growth.

Lastly, this study examined whether the *A. nidulans* SclB function is conserved in other ascomycetes species with different life styles that follow different strategies for colonizing their environments. Heterologous expression of Scl2 from the *V. dahliae* plant fungal pathogen, into *A. nidulans* Δ*sclB*, showed that its function seems to be conserved between ascomycetes. SclB/2 in the soli inhabitant *A. nidulans* as well as in the vascular pathogen *V. dahliae*, which is overwintering *ex planta* in the soil, seems to operate, as least partially in a conserved manner (**Fig. 6**). It will be interesting to examine in the future in more detail, to what degree the targets of *Vd* Scl2 and the *An* SclB are alike.

In summary, our data are presented in a model (**Fig 7**), which places SclB in a prominent position in a signaling pathway, where it orchestrates and promotes the passage of the fungus from the undifferentiated vegetative growth to the asexual development. SclB, ensures the activation of several mandatory regulatory circuits necessary for the onset of conidiation by its *in vivo* direct association with promoters of key regulatory genes, such as *brlA*, *ppoC* and *sclB*. Our work further establishes the SclB as a main actor to the scenery of central developmental pathway for the asexual sporulation. Asexual spore formation is an essential step for ascomycetes to spread and a better knowledge of the control of the fungal SclB regulatory function might contribute to a better control of fungal growth and dispersal in different environments.

**FIG 7:**
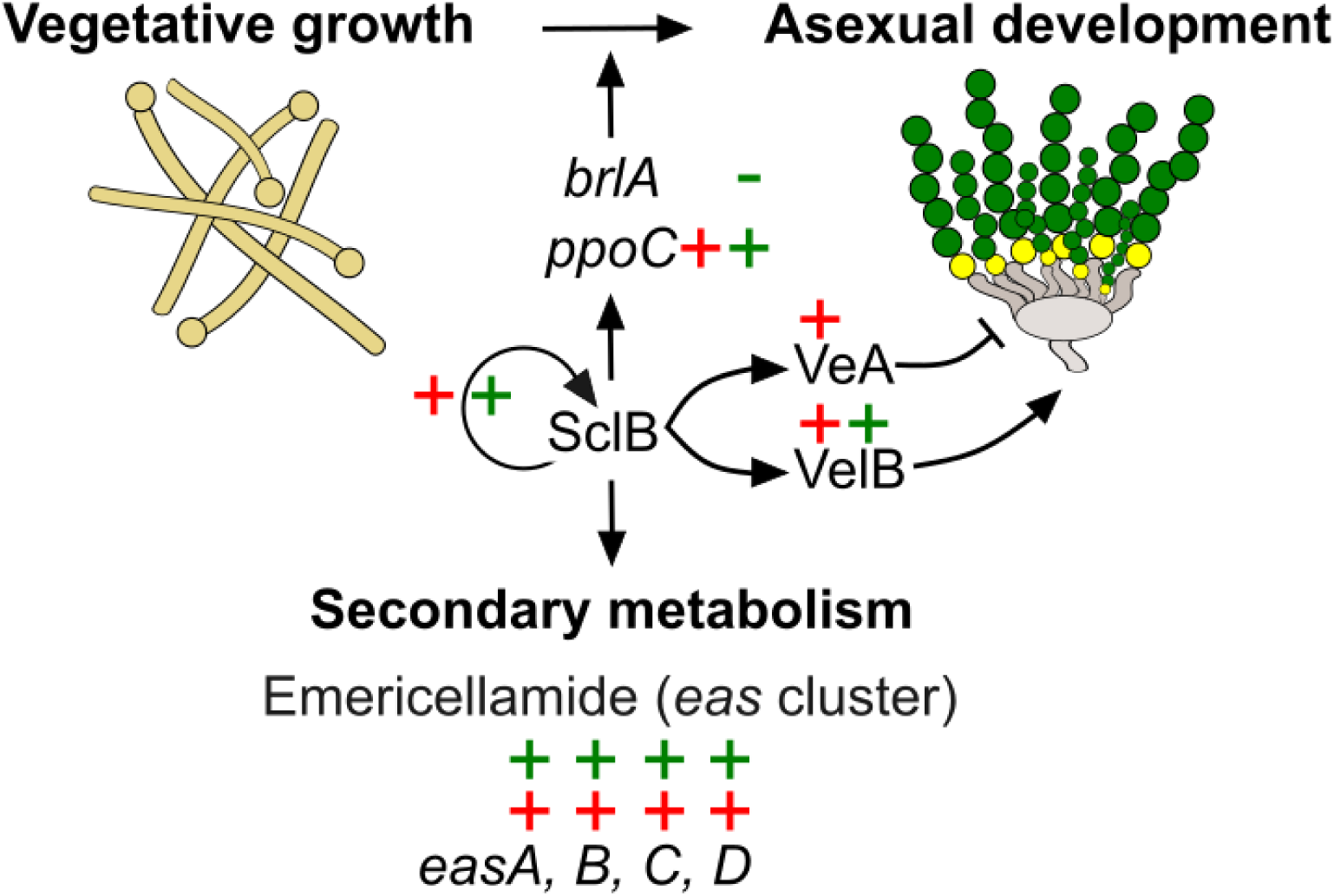
SclB is the organizer and coordinator for the transition of the fungus from the vegetative to the asexual growth. The graphic representation illustrates a current model where SclB orchestrates the shift of the fungus towards the asexual reproduction. The BrlA, PpoC, and SclB are established master regulators specifically of asexual development that are transcriptionally controlled directly by SclB during this transition. Additionally, SclB directly influences the transcription of *veA* and *velB* at the vegetative phase. However, the expression of *velB* (encoding for the VelB asexual inducer) but not of *veA* (encoding for the VeA asexual suppressor) is continuing to be induced shortly after the transition to the asexual phase. SclB is also associated *in vivo* to all four genes (*easA*, *B*, *C*, *D*) of the *eas* cluster, impacting in that manner directly the synthesis of emericellamides. The crosses, red for Veg and green for Asex conditions, just beside the genes, indicate an induction of gene expression based as derived from the generated ChIP-seq and RNA-seq data simultaneously. The *brlA* is the only gene that is repressed (indicated in the graph by a green hyphen) by SclB 3 h post induction of asexual growth. The designing of the graphical model was performed with the use of the freely available vector graphics software Inkscape (https://inkscape.org/).

## Material and Methods

### Strains, media and growth conditions

All the *E. coli*, *A. nidulans* and *V. dahliae* strains used in this study are listed in the **S1 Table**. As Wt strains *A. nidulans* AGB551 (32) and *V. dahliae* JR2 were used (33).

Minimal medium for *A. nidulans* had the composition: 1% (w/v) glucose, 2 mM MgSO_4_, 1x AspA (7 mM KCl, 70 mM NaNO_3_, 11.2 mM KH_2_PO_4_, pH 5.5), 0.1% (v/v) trace element solution (76 µM ZnSO_4_, 178 µM H_3_BO_4_, 25 µM MnCl_2_, 18 µM FeSO_4_, 7.1 µM CoCl_2_, 6.4 µM CuSO_4_, 6.2 µM Na_2_MoO_4_, 174 µM EDTA) and pH 5.5 (34). For solid medium 0.1% (w/v) uracil and 2 % agar were added. Liquid medium was supplemented with 5 mM uridine and 0.1% (v/v) pyridoxine. *A. nidulans* protoplasts were grown on solid medium supplemented with 120 mg/mL nourseothricin for selection. Selection-marker cassettes were recycled while growing the strains on medium with 0.5% (w/v) xylose/0.5% (w/v) glucose. Strains were grown at 37 °C.

*V. dahliae* strains were cultivated and transformed as described previously at (35).

Cloning was performed with *E. coli* DH5α. A lysogeny broth (LB) (36) was the liquid culture medium (1 % tryptone, 0.5 % yeast extract, 1 % NaCl, supplemented with 100 µg/mL ampicillin for selection) used for the propagation of *E. coli* cells transformed with plasmids according to (37). For solid medium 2% (w/v) agar was added. *E. coli* strains were grown at 37 °C.

### Plasmid manipulations and cloning

The backbone plasmid for construction of *A. nidulans* transformation cassettes was pME4696, initially used by Meister et al., (38). This vector carries a *Pml*I and a *Swa*I restriction site for integration of the 5’- and 3’- flanking regions of the target gene. These flanking parts are dictating the exact position in the *A. nidulans* genome where the cassette is integrated. Amplification of 5’- and 3’- regions was performed with AGB551 genomic DNA (gDNA). Assembly of the amplicons with pME4696 was performed by employing the GeneArt Seamless Cloning and Assembly Kit (Invitrogen, Carlsbad, CA, USA). All final cassettes carried nourseothricin resistance cassettes (natRM), as a recyclable marker. Isolation of plasmid DNA was conducted by the NucleoSpin Plasmid Kit (Macherey-Nagel, Düren, Germany) according to manufacturer’s instructions. Prior to excision of the cassette, all plasmids were verified via Sanger sequencing by Microsynth Seqlab GmbH (Göttingen, Germany).

### Assembly of the *Vd scl2::hinge::GFP::3’UTRAnsclB::trpC-*terminator cassette and the *Vd*-Scl2-GFP *A. nidulans* strain

The *Vd scl2* was initially amplified from gDNA template by using the primers RH835/836. The PCR product was subsequently cloned into pJet (pME5569) and sent for sequencing. The exact annotated sequence of the gene is presented into **S7 Fig.** It followed the junction of the *Vd scl2* with the linker, GFP (green fluorescent protein) and the marker. Fragment-1 consisted of a 5’-flanking region (amplified from gDNA template) and the gene (derived from the previous pJet vector) was constructed by using RH838/844. The fragment-2, composed by the linker, GFP and the marker, amplified from the plasmid pME5072 (39) with the primers AO165/ML8. The third fragment was generated by a 3’- flanking region, from gDNA template amplified with the primers RH840/841. All three fragments were subsequently cloned via a single Seamless reaction into the backbone vector pME4548 (40), previously digested by *Stu*I and *Eco*RV. The final plasmid was named as pME5567.

It followed the cloning of the cassette for the the strain *Vd* Scl2-GFP of *A. nidulans*. The 5’-flanking part was composed of four fragments: a 1898 bp amplicon spanning from the promoter of *An sclB* till the end of the genès *5’-UTR* (MB1569/1570), a 3234 bp amplicon for the *Vd scl2::hinge::GFP* (MB1571/1572), a 637 bp fragment starting from the start of the *3’-UTR* of the *An sclB* (MB1573/1574) and a 716 bp amplicon for a *trpC* terminator (MB1575/1576). The templates for each amplification were: for the first and the third part AGB551(Wt) gDNA, for the second was the plasmid pME5567 carried the coding sequence of *Vd scl2* from *V. dahliae* fused with a linker (hinge) and a GFP and for the fourth amplicon the plasmid pChS242 (41) was used carrying the sequence of the *trpC* terminator. The junctions of all parts constituting the whole 5’-flanking, took place by fusion PCR. When the 5 flanking part was prepared it was incorporated into the *Pml*I site of pME4696. The 1510 bp 3’-flanking region was amplified from gDNA of AGB551 with the primers (MB1577/1578), that was then introduced into the *Swa*I site of the same vector that carries the 5’-flanking part. The final plasmid after the incorporation of both flanking regions was named as pME5564 vector. Finally, the cassette was excised with the *Mss*I (*Pme*I) restriction enzyme and used subsequently for transformation in AGB551(Wt) and AGB1007(Δ*sclB*) *A. nidulans* strains leading to the corresponding strains AGB1711 and AGB1713, after the recycling of the selection marker.

### Assembly of the *Vd scl2::3’UTR AnsclB:trpC-*terminator cassette and the *Vd*-Scl2 *A. nidulans* strain

The construction of the of this cassette was performed in the same manner, using the same sets of primers, and even the same templates as used for the previous cassette of *Vd scl2::hinge::GFP::3’UTR_AnsclB_::trpC-*terminator. The only difference was the set of primers used to amplify the *Vd Scl2* from the corresponding vector that carried the coding sequence of *Vd scl2* fused with the *linker* and the *GFP* sequence. In that case it was used the set of primers MB1571/1579 (instead of MB1571/1572), where specifically the reverse primer (MB1579) ends the corresponding amplicon just after the stop codon of the *Vd scl2* and it is also designed to be fussed directly with the *3’UTR _AnsclB_* with a fusion PCR. Once the plasmid has incorporated its 5’- and 3’- flanking regions named as pME5565. The cassette was then excised and transformed into AGB551(Wt) and AGB1007(Δ*sclB*) *A. nidulans* strains, which after the after the recycling of the selection marker led to the generation of the AGB1715 and AGB1717 strains correspondingly.

### Assembly of the *ΔVd scl2* cassette and the *ΔVd scl2 V. dahliae* strain

The 5’ (1211 bp, primers RH838/RH839) and 3’flanking regions (1000 bp, primers RH840/RH841) were amplified from WT gDNA. The Nourseothricin resistance marker cassette (2194 bp, primers ML8/ML9) was amplified from pME4815 (42). The fragments were fused to pME4548, which was restricted with *Stu*I and *Eco*RV before using the GeneArt Seamless Cloning and Assembly Kit. The resulting plasmid pME5566 was transformed into the *V. dahliae* WT(JR2) strain and transformant VGB0685 was verified by Southern hybridization.

### Phenotypical assays

For examination of *A. nidulans* colonies, 2000 spores of each strain were spotted on MM plates. After 3-d of asexual growth, plates were scanned from top and bottom. *V. dahliae* phenotyping was performed by spotting 5x10^4^ spores of each strain on pectin-rich SXM; colonies were pictured after 5-d.

Quantification of *A. nidulans* conidia was performed with fresh spores from which identical number were distributed equally on MM plates. After 3-d of asexual growth, the spores were collected and quantified. The quantification of *V. dahliae* spores was done as described before Sasse et al., (41). The spores of both fungi were counted with the Coulter Z2 particle counter (Beckman Coulter GmbH, Krefeld, Germany).

### ChIP-seq

The GFP-SclB (AGB1010) strain of *A. nidulans*, expressing *sclB* fussed in its N’-terminus the GFP under its native promoter, was used to perform ChIP-seq experiments under Veg and Asex conditions. A total number 5x10^8^ spores was inoculated in 500 mL liquid medium inside of 2 L flasks and grown for 20 hours under constant rotation, light and at 37 °C. For the Veg ChIP, after the 20 h incubation, mycelia were dried and immersed in 1 % formaldehyde fixing solution for 20 minutes. For the 2^nd^ ChIP, mycelia were transferred after the 20 h vegetative growth to MM plates and grown asexually for additional 3 h. Then fixation was performed as previously described. For the Veg ChIP three biological replicates for GFP-SclB and the Wt strain (negative control/background signal) were used. For the Asex ChIP, were used three biological replicates for GFP-SclB and two for the Wt strain; in the subsequent analysis for the identification of the peaks, the sequenced data for the second biological replicate of the Wt were used as a control for the second and the third biological replicate of the GFP-SclB sequencing data. The GFP antibody (Abcam ab290) was applied to the IPs. The rest of the ChIP protocol, library preparation, NGS sequencing and ChIP-seq data analysis was performed as describe in Sasse et al., (41). ChIP-seq library constructions and sequencing of the ChIP samples was performed in NGS-Integrative Genomics Core Unit (NIG), University Medical Center Göttingen.

The pipeline for the ChIP-seq analysis was as described by Sasse et al., (41). Web tools for the analysis were provided either from the GALAXY or the RStudio platforms both maintained by the GWDG (Gesellschaft für wissenschaftliche Datenverarbeitung mbH Göttingen). For the mapping of the raw sequences, the *A. nidulans* genome (FungiDB-46_AnidulansFGSCA4_Genome.fasta) was used. The raw sequencing data for the ChIP-seqs [BioProject ID PRJNA1292721] and for the RNA-seqs [BioProject ID PRJNA1293339] have been deposited at NCBI.

### DNA and RNA extraction

Genomic DNA was extracted from mycelia grown overnight in light and at 37 °C. DNA isolation was described by Thieme et al., (18).

Liquid cultures were inoculated with 10^8^ spores in 100 mL medium. After 20 h growth with constant agitation in light, vegetative mycelia were dried and 100 mg for each sample were snap frozen in liquid nitrogen. For the Asex RNA-seq, mycelia were grown vegetatively for 20 h and then transferred to MM plates for another 3 h. Mycelia were processed as described before. The extraction of RNAs was done as described in Bastakis et al., (28).

### Southern hybridization

The positive clones, derived from the transformations in *A. nidulans* and *V. dahliae* genome, were further checked genetically, for integration of the cassette, by Southern hybridization as described by Southern, 1975 (43). The labelling of the Southerns’ probes was done by AlkPhos Direct Labelling Module (GE Healthcare Life Technologies, Little Chalfont, UK) following the manufacturer’s instruction.

### RNA-seq and data analysis

The preparation of the RNA-seq libraries and the following sequencing, from the step of the quality’s check of the initial RNA samples till the quality of sequencing followed the pipeline as described by Szemes et al., (44).

The RNA-seq analysis was performed in GALAXY platform (45). as provided by GWDG. Raw sequencing reads were mapped in the *A. nidulans* genome (downloaded from fungidb.org: FungiDB-46_AnidulansFGSCA4_Genome.fasta) by using the Bowtie2 (46) (Galaxy Version 2.3.4.2). Subsequently, matrices were prepared, by employing the htseq-count tool (Galaxy version 0.9.1), that were then used to calculate the DEGs of the deletion Δ*sclB* versus the Wt samples, with the Galaxy implemented tool DESeq2 (47) (Galaxy version 2.11.40.6+galaxy2).

### Microscopy

Fluorescence microscopy was performed as described in Bastakis et al., (28). The nuclear localization of the *Vd* Scl2-GFP was accessed (including the Wt/negative control), by the expression of RFP-H2A. The integration of the gene encoding the nuclear marker was performed into the corresponding *A. nidulans* strains by the transformation of the cassette ^p^*gpdA::intron::mrfp::h2A*(*cDNA*) carried by plasmid pME3173 (48), resulting into the strains, AGB1718, AGB1719 an AGB1720.

### Figures processing

The processing of the all figures was done by the vector-graphics editor Inscape (Inkscape Project, 2020; Inkscape, available at https://inkscape.org).

## Acknowledgments

We would like to thank Dr. Gabriela Salinas and Fabian Ludewig from the NGS-Integrative Genomics Core Unit (NIG), University Medical Center Göttingen for their excellent assistant on our NGS-based approaches.

## Funding

Two grants from the Deutsche Forschungsgemeinschaft (DFG) financially supported this work: grant BR1502/15-2 and grant IRTG PRoTECT, both of them awarded to Gerhard H. Braus. We acknowledge the support by the Open Access Publication Funds of the Göttingen University. The funders had no role in study design, data collection and analysis, decision to publish, or preparation of the manuscript.

## Supplementary material

**FIG S1:**
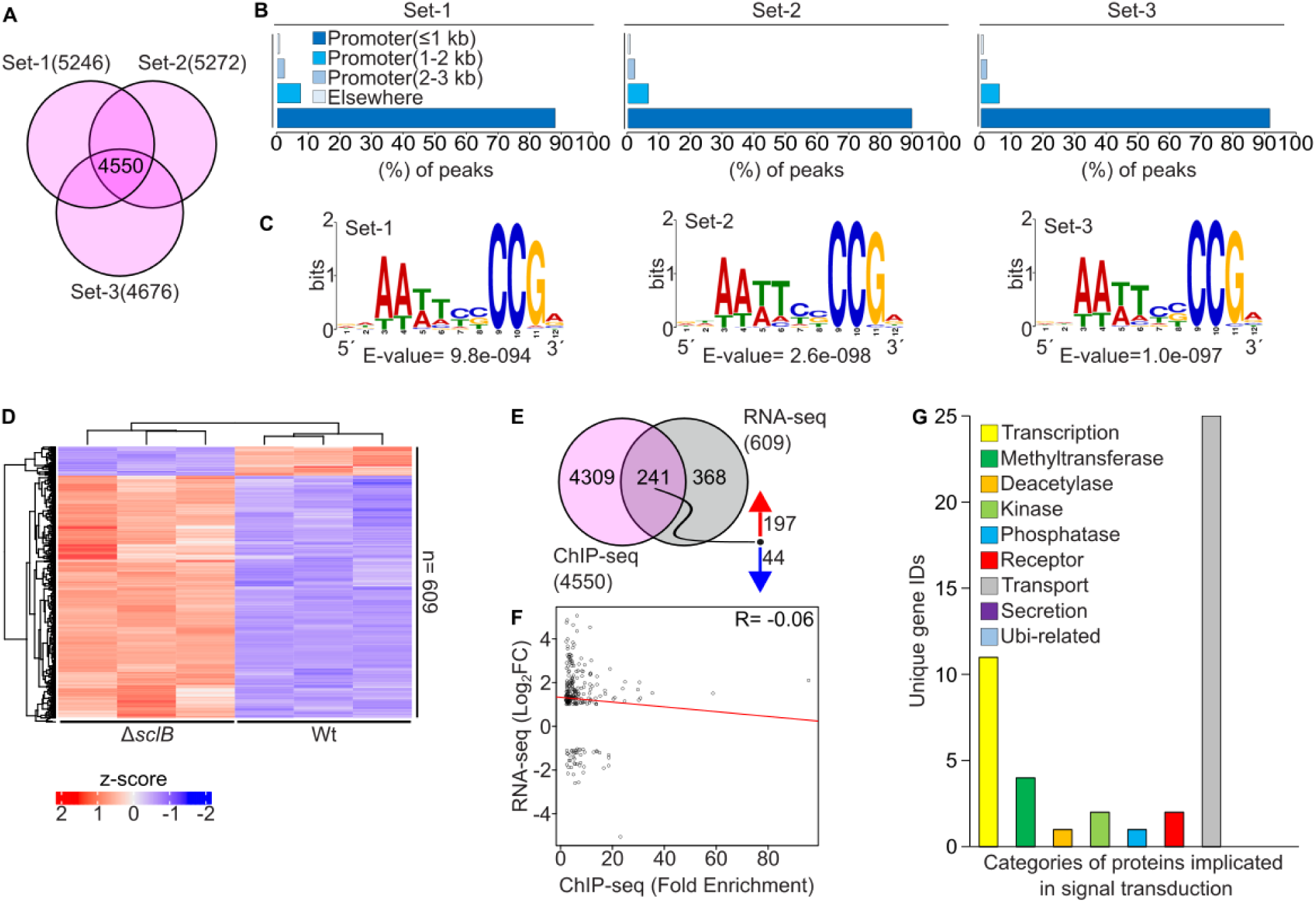
*In vivo* binding landscape of the GFP-SclB transcriptional regulator during early *A. nidulans* asexual development. (**A**) The Venn diagram presents the overlap of three independent sets of ChIP-seq analysis for GFP-SclB versus Wt. ChIP-seq was performed in light under Asex growth conditions. In sum, 4550 unique gene locus IDs, found to have peaks (cut offs: p < 0.05 and fold enrichment (F.E.) ≥ 2.0) located in up to 3 kb promoter regions, discovered at the same time in all three independent sets of the ChIP-seq samples. (**B**) Bar diagrams depict the distribution of the statistically significant ChIP-seq peaks over different genetic structures for each of the three independent sets of this analysis. For all three sets around 90 % of the discovered peaks are located within promoter regions of genes up to 1 kb from the transcriptional start site. (**C**) Top scored logos, as discovered by performing a *de novo* motif analysis by employing the MEME-ChIP tool. For each independent set of analysis, it was used as an input a set of 150 sequences, each with 100 bp length. All of those sequences were located underneath the summits of the top-ranked 150 ChIP-seq peaks for each set. The summits of all of these peaks were located up to 3 kb promoter regions. (**D)** Heatmap is illustrating differential expression of genes (as z-score values) from mRNAs of Δ*sclB* and Wt strains, derived from Veg growing mycelia. The number n=609 represents the total number of genes that were differentially expressed under the cut offs: p < 0.05 and -1 ≤ log_2_FC(Fold Change) ≥ 1. The RNA-seq was performed with three independent biological replicates of the Δ*sclB* and for the Wt strain as well. (**E**) Venn diagram presenting the overlap among the highly reproducible targets genes from the ChIP-seq with GFP-SclB (**A**) with the DEG from the RNA-seq of Δ*sclB* (**B**). Both experiments were performed with mycelia Asex growth conditions. The corresponding cut offs were for the ChIP-seq: p < 0.05 and fold enrichment (F.E.) ≥ 2.0 and for the RNA-seq: p < 0.05 and -1 ≤ log_2_FC(Fold Change) ≥ 1. The numbers at the right side of the red and blue arrows indicate the total number of genes that their expression was increased or decreased, from the total genes of the overlap. The set of these 241 gene locus IDs is characterized as the direct *in vivo* targets of GFP-SclB early during the asexual development. (**F**) Pearson correlation coefficient, presented by a scatter plot, among the peak’s fold enrichment from the ChIP-seq performed under Asex growth conditions with the gene expression (as of Log_2_FC) from the RNA-seq, under the same growth conditions. For this plot only the 441 direct target genes as discovered by the overlap of ChIP-seq with the RNA-seq in (**E**) were used. The R= -0.06 at the upper right corner of the plot depicts the negative correlation among the compared NGS data sets that were examined. (**G**) Bar chart showing the number of genes from the overlap (241 gene locus IDs) between the ChIP-seq and RNA-seq (as showed previously in **E**) belonging to categories of genes encoding for proteins, known to play crucial roles in signal transduction pathways. The number for each category assigned after conducting manual examination in the set of 241 gene locus IDs for each indicative protein category of the signaling.

**FIG S2:**
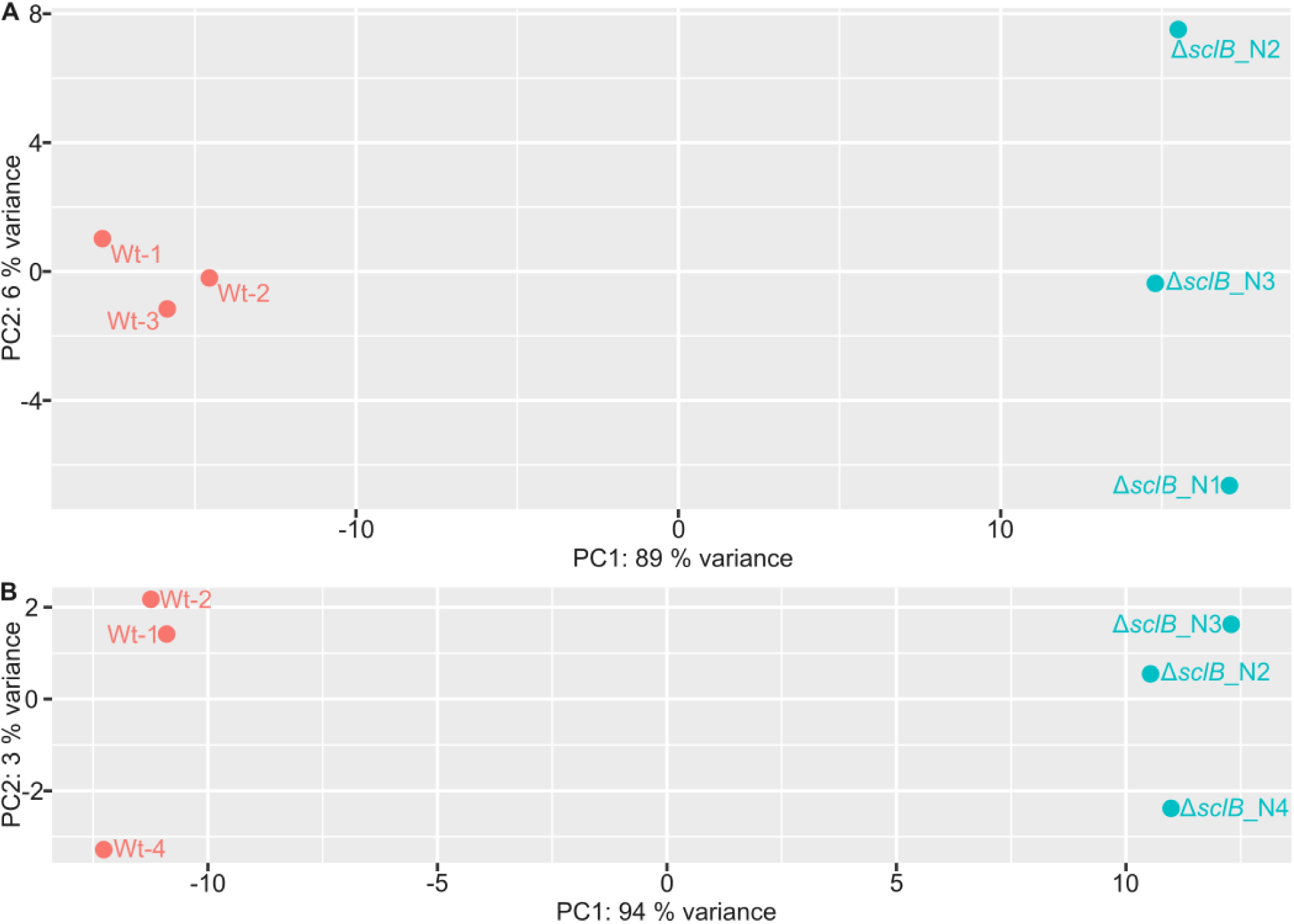
Clustering of different samples for the RNA-seq performed during *A. nidulans* vegetative growth or induction of asexual spore formation for Wt compared to Δ*sclB* mutant strains. Principal component analysis (PCA) among the RNA-seq samples of Wt and Δ*sclB* either growing under (**A**) Veg or (**B**) Asex conditions.

**FIG S3:**
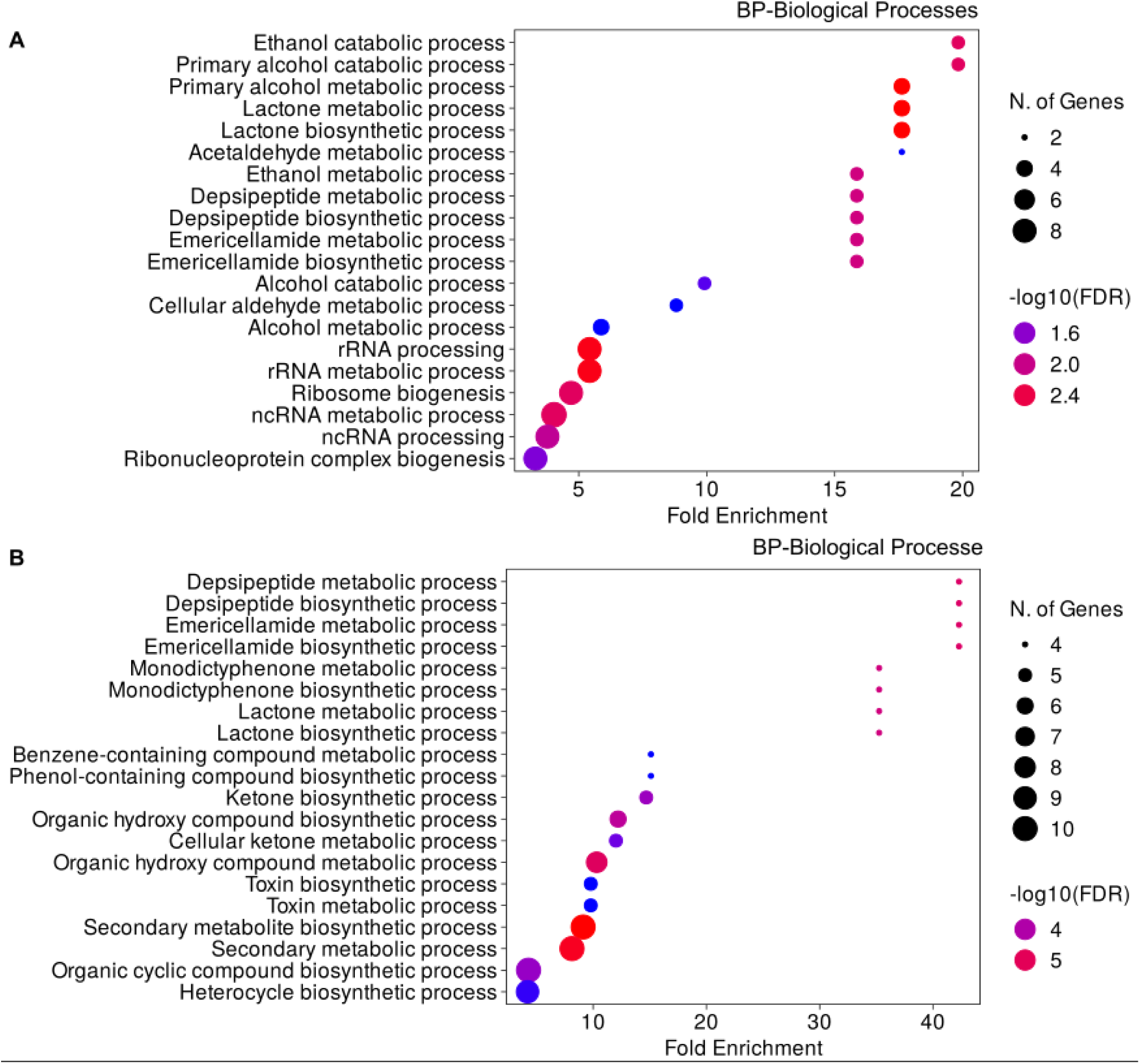
Proteins involved in several *A. nidulans* metabolic processes are among the top direct target genes of SclB either during vegetative growth or when asexual development is induced. Bubble plots presenting the Gene Ontology (GO)-enrichment analysis (in terms of biological processes/BP), of genes belonging to the overlap of ChIP-seq with the RNA-seq for mycelia either growing (**A**) under Veg (441 locus IDs) and (**B**) Asex (241 locus IDs), conditions respectivelly. All IDs were submitted to cut offs for the ChIP-seq: p < 0.05 and fold enrichment (F.E.) ≥ 2.0 and for the RNA-seq: p < 0.05 and - 1 ≤ log_2_FC(Fold Change) ≥ 1, prior of the enrichment analysis. The coloring of the bubbles corresponds to statistical differences presented as -log10(FDR). The size of the black circles reflects the size of gene sets per different identified GO-categories of biological process (BP), under which the GO-enrichment analysis was performed.

**FIG S4:**
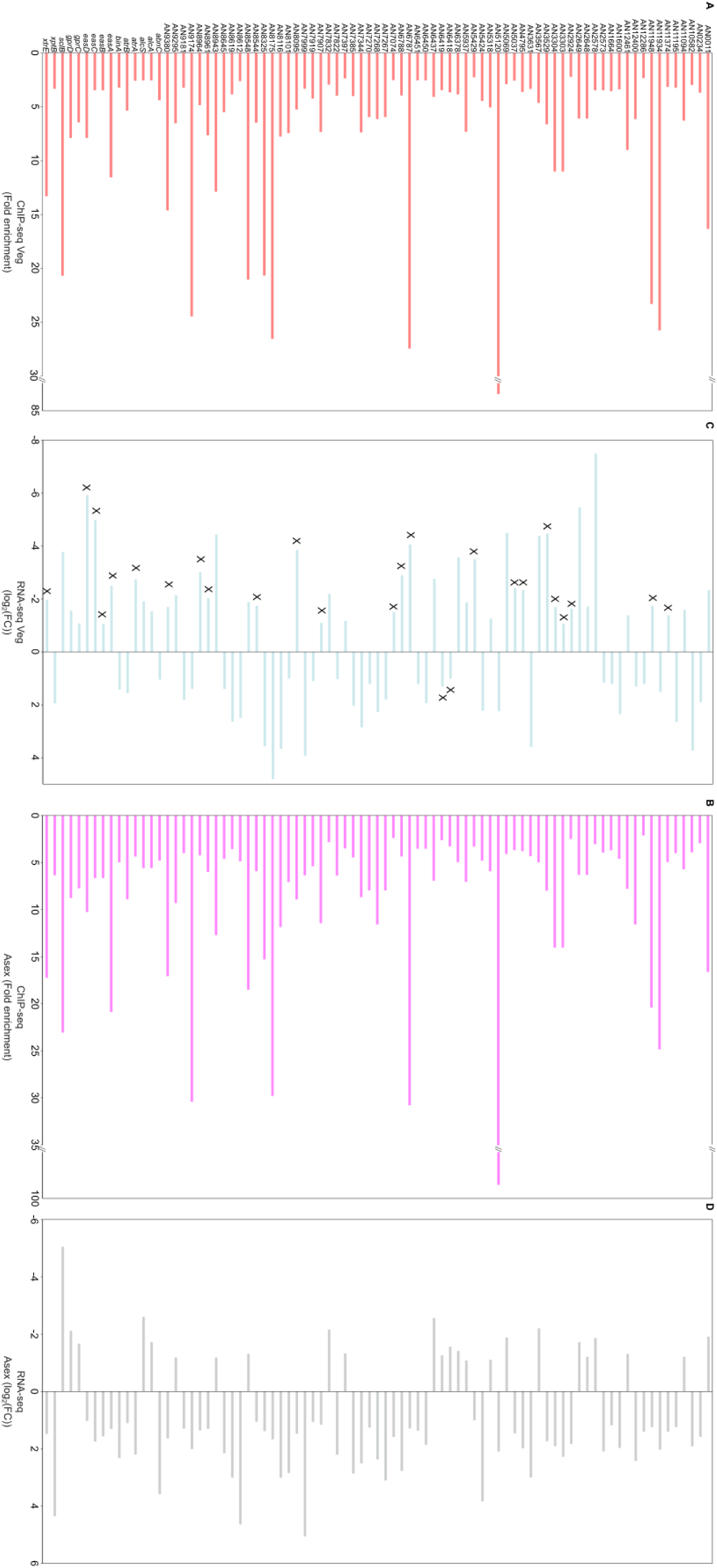
Specific set of common SclB controlled genes during vegetative growth or asexual development of *A. nidulans*. Bar diagrams illustrating (**A** and **C**) the ChIP-seq results (fold enrichment) and (**B** and **D**) the RNA-seq results (log_2_FC), from mycelia grown under Veg or Asex conditions. Each diagram presents the corresponding values for each of the 83 gene locus IDs that were found to be simultaneous targets of GFP-SclB (in both ChIP-seqs data sets) and differentially expressed (in both RNA-seqs data sets), independently from the growth conditions. The applied thresholds for the ChIP-seq were: p < 0.05 and fold enrichment (F.E.) ≥ 2.0, the peak must be located up to 3 kb upstream from the TSS of every gene locus; for the RNA-seq: p < 0.05 and -1 ≤ log_2_FC(Fold Change) ≥ 1. The letter X in (**C**) bar plot, indicates the genes that show contrary expression pattern compared to one during the Asex growth (**D**).

**FIG S5:**
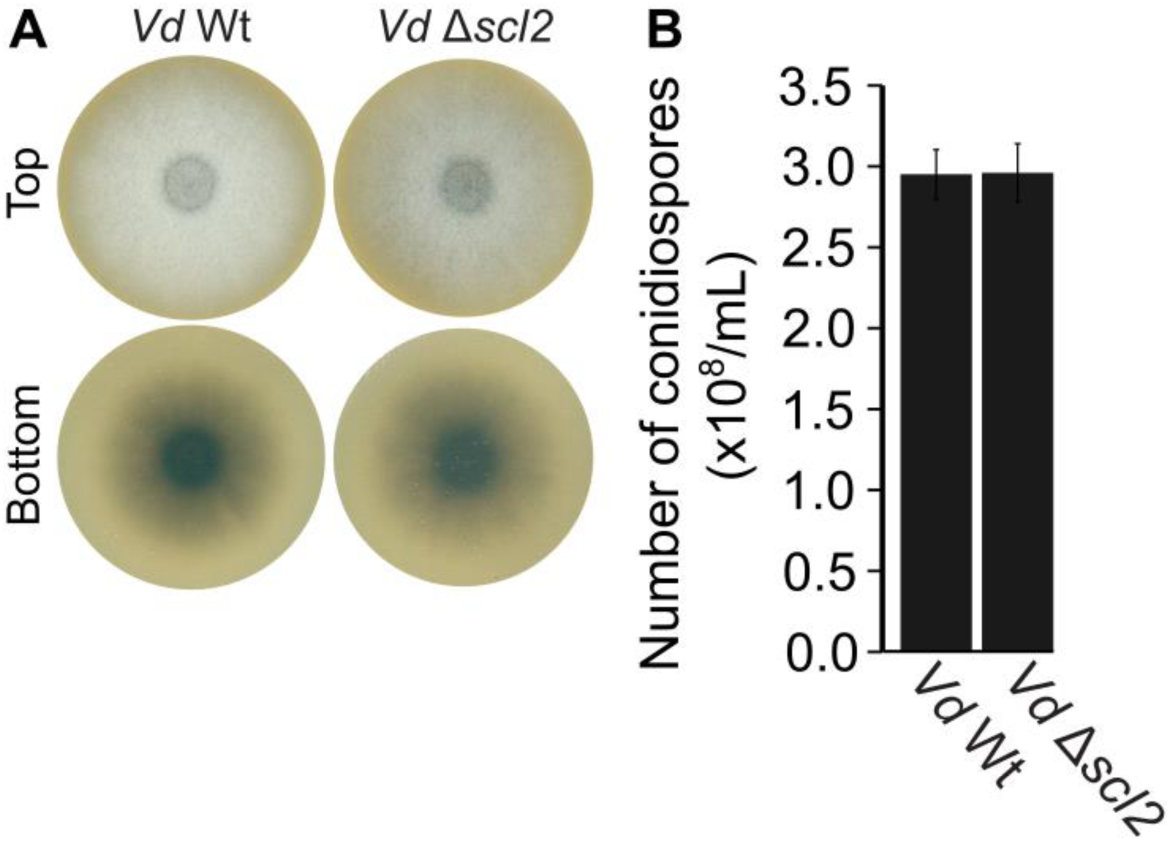
*V. dahliae* Scl2 is dispensable for colony formation and conidiation. (**A**) Colony growth of the *V. dahliae* Wt-JR2 in comparison to two independent transformants of the *scl2* deletion strain on pectin-rich simulated xylem medium (SXM). (**B**) Quantification of conidiospores produced by either the Wt or 2 independent *scl2* deletion strains in liquid SXM culture after 5 days of incubation. Bars represent the mean values of two independent experiments with 3-4 repetitions each.

**FIG S6:**
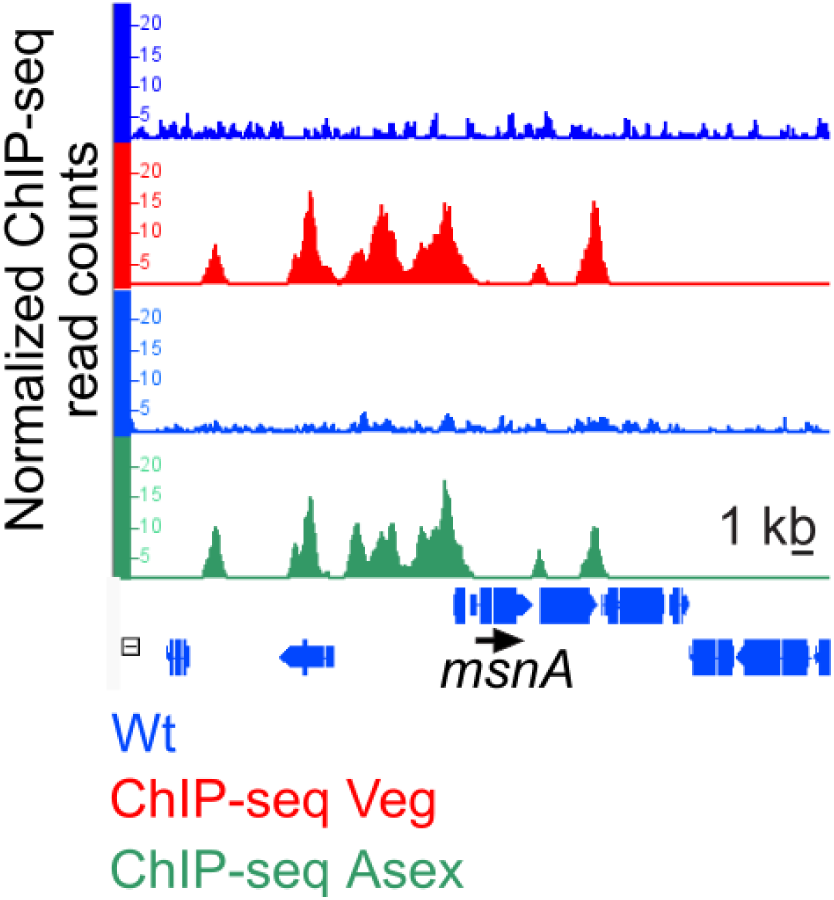
SclB is associated *in vivo*, in multiple sites of the promoter of *msnA*. The snapshot from the IGB showing the *in vivo* binding of SclB (ChIP-seq peaks) on the promoter of *msnA gene*. Peaks appeared in the red tracks referred to the ChIP-seq performed under Veg conditions and peaks showed in green tracks under Asex conditions correspondingly. Blue tracks, in each case, referred to negative control (background signal) of each ChIP-seq.

**FIG S7:**
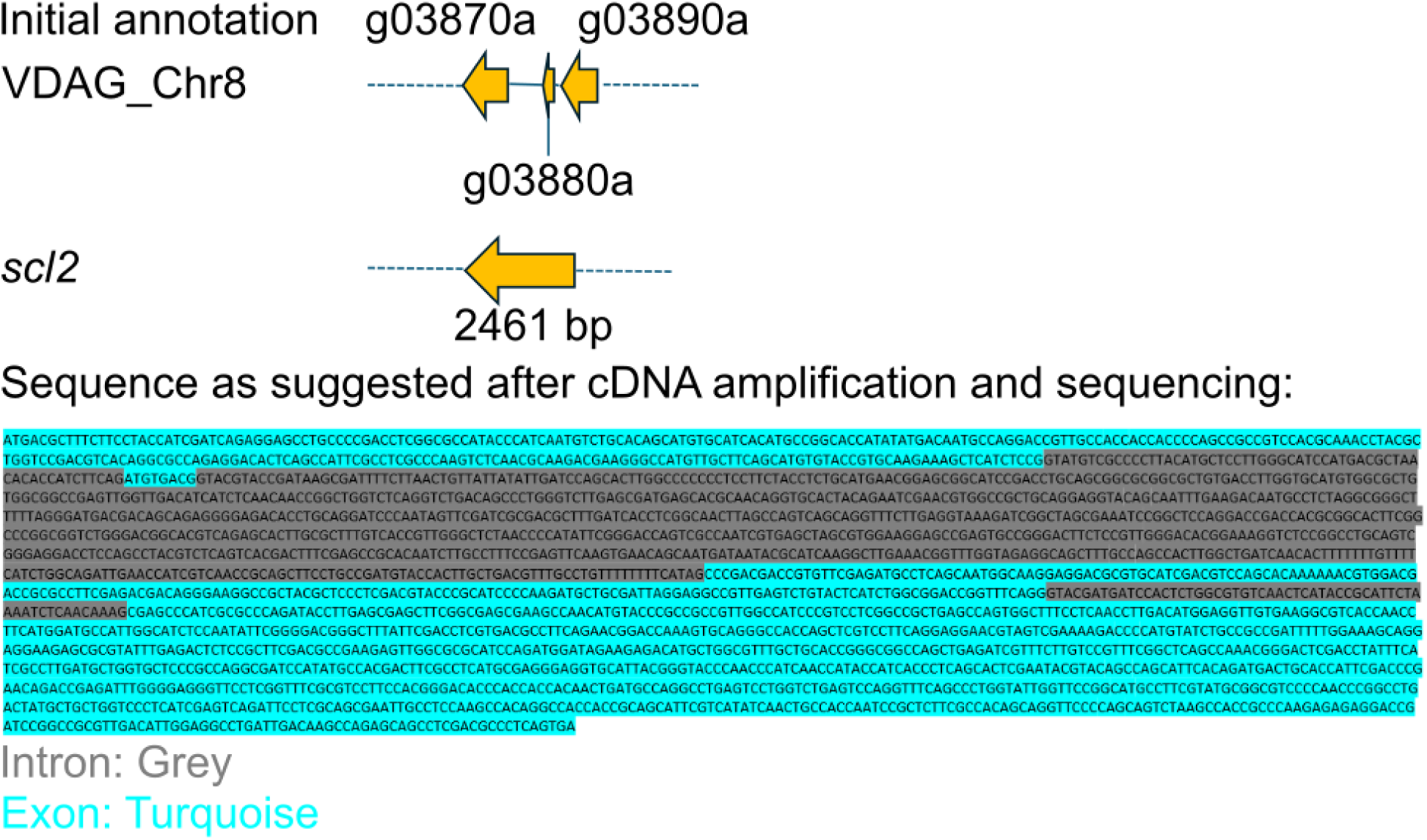
Prediction and annotation of the *scl2* in *V. dahliae*.

**S1 Table:** Primers used in this study.

**S2 Table:** Strains (*A. nidulans*, *V. dahliae* and *E. coli*) used in this study.

**S3 Table:** Target genes from the ChIP-seq experiments and differentially expressed genes from the RNA-seq experiments.

